# Reduced antigenicity of Omicron lowers host serologic response

**DOI:** 10.1101/2022.02.15.480546

**Authors:** Jérôme Tubiana, Yufei Xiang, Li Fan, Haim J. Wolfson, Kong Chen, Dina Schneidman-Duhovny, Yi Shi

## Abstract

SARS-CoV-2 Omicron variant of concern (VOC) contains fifteen mutations on the receptor binding domain (RBD), evading most neutralizing antibodies from vaccinated sera. Emerging evidence suggests that Omicron breakthrough cases are associated with substantially lower antibody titers than other VOC cases. However, the mechanism remains unclear. Here, using a novel geometric deep-learning model, we discovered that the antigenic profile of Omicron RBD is distinct from the prior VOCs, featuring reduced antigenicity in its remodeled receptor binding sites (RBS). To substantiate our deep-learning prediction, we immunized mice with different recombinant RBD variants and found that the Omicron’s extensive mutations can lead to a drastically attenuated serologic response with limited neutralizing activity *in vivo*, while the T cell response remains potent. Analyses of serum cross-reactivity and competitive ELISA with epitope-specific nanobodies revealed that the antibody response to Omicron was reduced across RBD epitopes, including both the variable RBS and epitopes without any known VOC mutations. Moreover, computational modeling confirmed that the RBS is highly versatile with a capacity to further decrease antigenicity while retaining efficient receptor binding. Longitudinal analysis showed that this evolutionary trend of decrease in antigenicity was also found in hCoV229E, a common cold coronavirus that has been circulating in humans for decades. Thus, our study provided unprecedented insights into the reduced antibody titers associated with Omicron infection, revealed a possible trajectory of future viral evolution and may inform the vaccine development against future outbreaks.

## Introduction

The severe acute respiratory syndrome coronavirus 2 (SARS-CoV-2) continues to evolve, producing variants of concern (VOC) with improved transmissibility and abilities to evade host immunity. The newly identified VOC Omicron (B.1.1.529) contains a large number of mutations including 11 that localize on the variable RBS, which is the major target of serologic response^1^. These mutations collectively facilitate the immune evasion of both vaccinated and convalescent sera^2–8^. However, it remains unclear if the extensive RBD mutations could affect the magnitude and immunodominance hierarchy of the host antibody response^9^.

## Results

### Deep learning predicts a decrease in the Omicron RBS antigenicity

Here, we used a geometric deep learning model (ScanNet^10^) for the prediction of epitopes to systematically investigate the RBD antigenic profiles for WT (the Wuhan strain) and the VOCs (**Figure 1**). To evaluate the accuracy of ScanNet, we calculated antibody binding propensity and found that it correlates well with the frequency of structurally determined RBD epitopes (Spearman rho=0.77, **Methods, Figure S1**). As expected, the RBS residues have high antibody binding propensity for all the RBDs (**Figure 1A**). Alpha, Beta, and Delta VOCs have a moderate increase in the RBS antigenicity compared to WT (**Figure 1B,E**). However, the antigenicity of Omicron RBS (particularly, residues 470-500 and 445-455) was significantly reduced (**Figure 1B-E**). Moderate increases in antigenicity were also detected for several residues (403-420, 501-505), however, these sites barely overlap with the dominant epitopes mapped experimentally (**Figure S2**). Together, our analysis indicates that the overall antigenicity of Omicron RBS is reduced.

**Figure 1.**
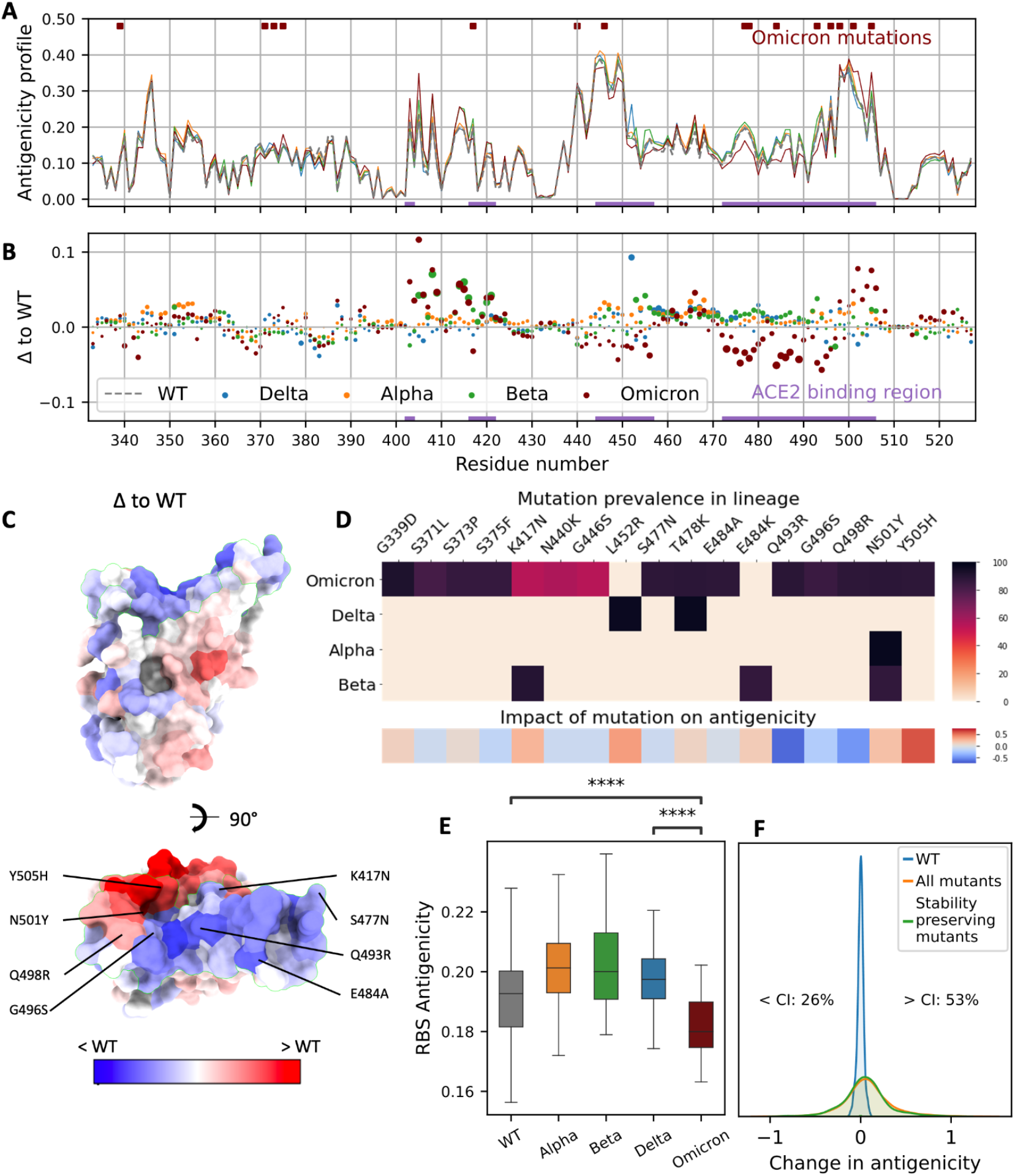
Impact of Omicron mutations on antigenicity based on Geometric Deep Learning. **A**. Residue-wise antigenicity profile of WT and four VOCs computed with ScanNet. **B**. Difference between each VOC and WT, depicted as a scatter plot. The area of each point represents the statistical significance of the difference (larger is more significant): it is proportional to the absolute value of the associated Z-score (clipped at |Z|=10, the dots in the caption correspond to |Z|=5). **C**. Omicron RBD colored by the difference of antigenicity (PDB: 7qnw) with respect to WT. **D**. Upper panel: Prevalence of mutations for each VOC based on GISAID. Bottom panel: Corresponding predicted change in overall antigenicity. **E**. Boxplots of RBS average antigenicity for WT and four VOCs calculated over multiple structures. Center line: median; box limits: upper and lower quartiles; whiskers: 1.5x interquartile range; p-value annotated legend: ns: p> 5e-2, ***: p<1e-4 (two-sided Wilcoxon-Mann-Whitney test). **F**. Distribution of changes in antigenicity across all single-point mutations and all stability-preserving single-point mutations previously identified by deep mutation scan (^22^, cut-off of −0.5 in log-odds scale). The blue histogram denotes the distribution over structural models of the WT scores, and intuitively corresponds to the noise level induced by the structural modeling component of the prediction pipeline. The corresponding matrix is shown in **Figure S3A**.

To assess the significance of the change and dissect the individual contribution of the Omicron mutations to the overall antigenicity, we modeled the structures of 15 single point mutants and calculated their antigenic profiles (**Methods**). Eight mutations (53%) decreased the antigenicity, in particular Q493R, G496S, and Q498R (**Figure 1D**). Five mutations (33%) increased the antigenicity while the remaining had no obvious effect. Next, we modeled the structures of all the point mutants and calculated their antigenicity (**Figure S3A)**. Only 26% decreased the antigenicity (**Figure 1F**). Therefore, the reduced antigenicity of Omicron is not random (p=0.034) and may result from evolutionary pressure.

### Omicron mutations lead to a drastic and systemic reduction in RBD antigenicity *in vivo*

To substantiate the deep-learning analysis, we immunized mice via the mucosal delivery route with the recombinant RBDs from WT (n=4) or VOCs (n=5) and analyzed their adaptive immune responses (**Methods**). All the animals showed robust and comparable T cell responses as indicated by the *in vitro* recall assays. Specifically, their splenocytes produced high levels of IFNg when re-stimulated with WT, Delta, or Omicron RBDs regardless of the immunogens that they originally received (**Figure S4A**), suggesting a successful initiation of Th1-mediated immune response. A strong Th17 response was also generated as expected for this type of mucosal immunization regimen^11^ (**Figure S4B**). IL-17 levels appeared to be more consistent among all groups of mice suggesting they were mainly produced by antigen-specific CD4 T cells whereas IFNg can come from nature killer or gamma-delta T cells without the need of antigen recognition. We also analyzed the local response in the lungs in the animals and observed comparable IFNg and IL-17 responses (**Figure S4C-D**).

Next, we performed ELISA to measure antibody titers of the immunized sera against the corresponding antigens. In contrast to the T cell response, we found that the antibody titers (half-maximal inhibitory reciprocal serum dilution or ID50) of the Omicron-immunized sera were significantly reduced by over 15-fold (mean ID50 = 924) compared to that of WT (mean ID50 = 15,325) and other VOCs (mean ID50s = 11,564, 14,683 and 19,557 for Alpha, Beta, and Delta, respectively) (**Figure 2A**). Thus, our *in vivo* experiments were consistent with the deep learning model, revealing that mutations can greatly reduce the antigenicity of Omicron RBD.

**Figure 2:**
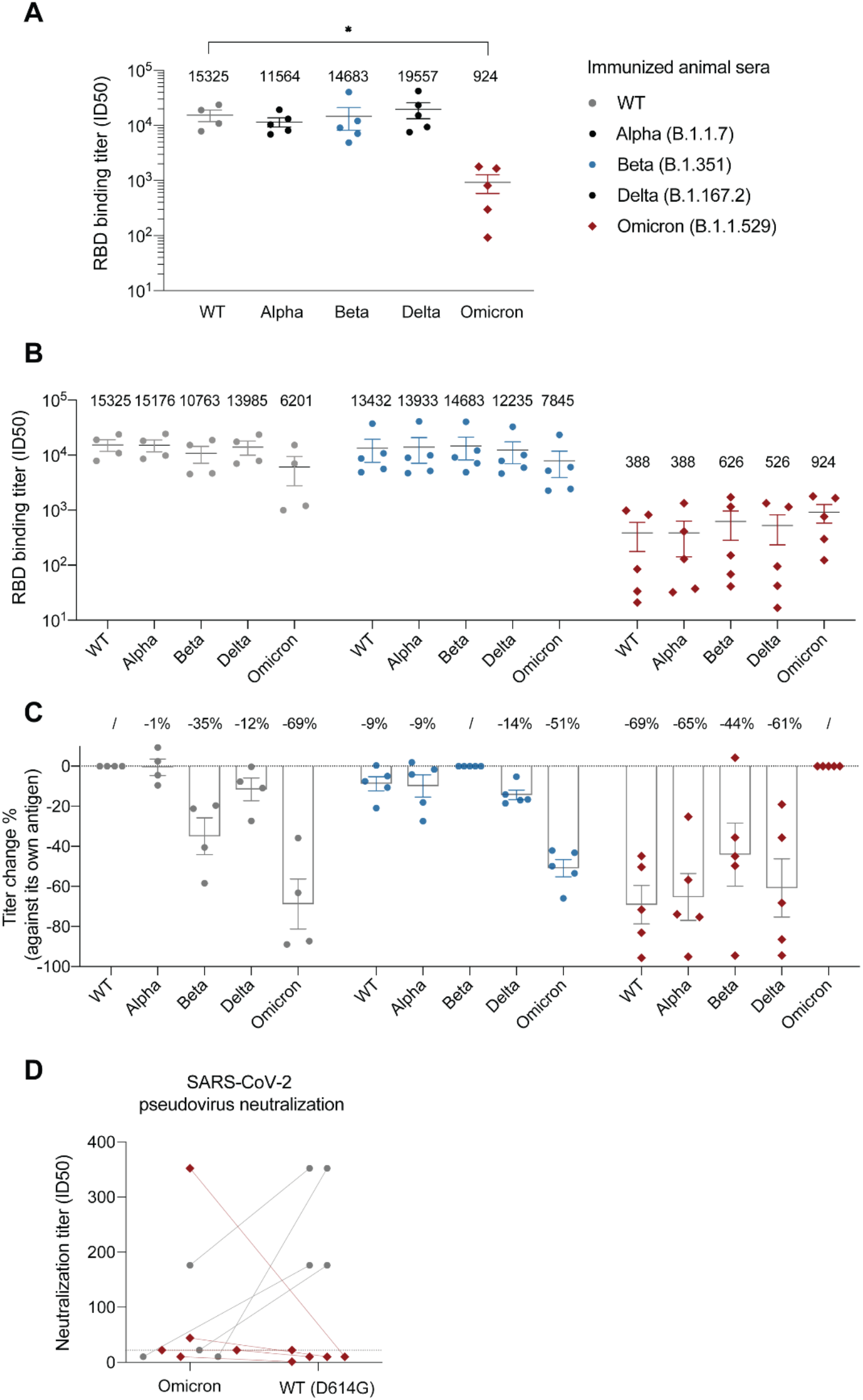
Analysis of the RBD-immunized sera. **A**. ELISA of RBD-immunized mouse sera against the corresponding antigen. Binding titer was calculated as the ID_50_ (reciprocal serum dilution that inhibits the 50% maximal RBD binding). **B**. ELISA of RBD-immunized sera against five different RBDs (cross-reactivity analysis). **C**. The percentage change of binding titers against different RBDs. **D**. Pseudovirus neutralization assay evaluating the potencies of WT and Omicron RBD-immunized sera against either SARS-CoV-2 WT (Wuhan-Hu-1, D614G) strain or Omicron. The neutralization titer was calculated as the ID_50_ (reciprocal serum dilution that inhibits the 50% of the maximal pseudovirus infection). Two connected dots referred to the pseudovirus neutralization results of the same animal serum. Dashed line indicates the highest serum concentration (i.e., dilution of 22 which is the lowest reciprocal serum dilution) used in the study.

Previous structural analysis reveals that the majority of antibodies target the variable RBS^12^. The remaining antibodies bind conserved epitopes that are cross-reactive among VOCs^13–16^. To better understand the antigenicity and immunodominance hierarchy of RBD variants, we evaluated the cross-reactivity of immunized sera by ELISA (**Figure 2B-C**). WT-immunized sera had comparably high titers against Alpha (ID50 = 15,176) and Delta (ID50 = 13,985) RBDs but presented decreased activities against Beta (ID50 = 10,763; by 35%) and more substantially against Omicron (ID50 = 6,201; by 69%). The magnitudes of antibody evasion by VOCs were consistent with clinical data^3–8^, indicating that the RBD immunodominance hierarchy is similar between mouse and human.

We found that Omicron-immunized sera had substantially lower antibody titers against all the VOCs (with the mean ID50s in the range of 388-626, **Figure 2B**). Despite the reductions, Omicron-immunized sera still bind most efficiently to its own antigen (**Figure 2B**), indicating that Omicron’s RBS remains to be highly antigenic while other conserved epitopes can also contribute to the overall antigenicity. Moreover, while the titer of Omicron-immunized sera against WT RBD was only a small fraction of that of WT-immunized sera against Omicron (388/6201 or 6%), the percentages of cross-reactive antibodies were highly comparable (~31%, **Figure 2C**). Thus, the immunodominance hierarchy for Omicron remained largely unaltered and the reduction of response was rather systemic, contributed by both RBS and other conserved epitopes. This result was further supported by competitive ELISA using either the recombinant ACE2 or high-affinity nanobodies targeting distinct and highly conserved RBD epitopes (**Figure S5, Methods**).

Since Beta RBD shares three mutation sites with Omicron (K417N, E484K/A, and N501Y) critical for antibody binding, we also evaluated the cross-reactivity of Beta-immunized sera and found that these sera cross-reacted better (49%) with Omicron than the WT sera (31%) (**Figure 2C**). Since the antibody titers of the Beta immunized mice are comparable to those of WT sera, we conclude that these three mutated residues do not significantly contribute to the antigenicity decrease (**Figure 1D**).

Next, we performed SARS-CoV-2 pseudovirus assay to evaluate the contribution of Omicron mutations to the neutralization potency of the immunized sera (**Figure 2D**). Despite some cross-reactivity of WT-immunized sera against Omicron (ID50 = 6,201), their neutralization activities were barely detectable. Strikingly, the potencies of the Omicron-immunized sera were generally inefficient against the Omicron virus (except for one serum) and their activities against WT (the Wuhan-Hu-1/D614G strain) were hardly detected.

### Analysis of the evolution of hCoV229E reveals a decrease in antigenicity

hCoV229E is a common cold coronavirus that has been circulating in human populations for decades. As one of the first coronavirus strains being described, its sequences and structures have been well documented and can be used as a model system to study the evolution of antigenicity and host serologic response^17–19^. Here, structural modeling followed by longitudinal analysis of hCOV229E RBDs revealed an overall trend of decreasing antigenicity on RBS until 2010s with subsequent oscillation during the last decade (**Figure 3A**).

**Figure 3:**
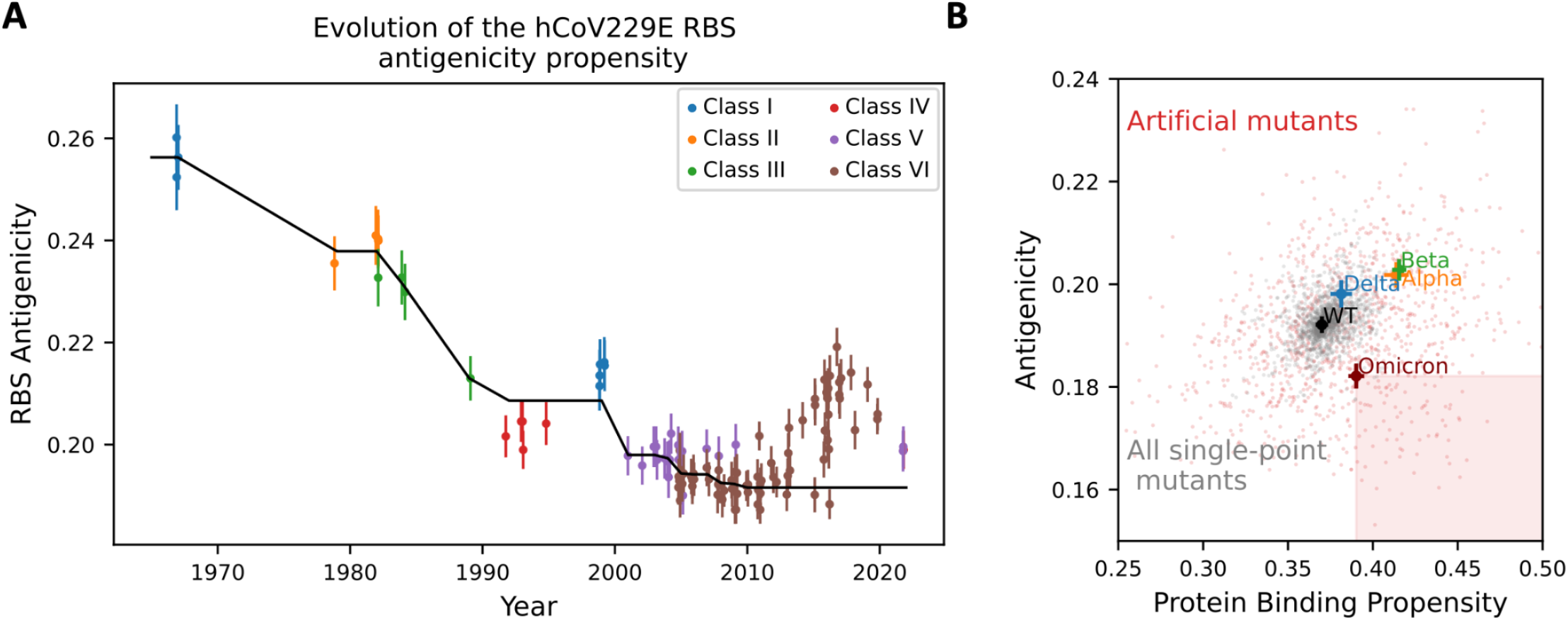
Plausibility of further decrease of antigenicity in future variants. **A**. Evolution of the antigenicity of hCoV229E RBS for isolates collected from the 1960s to date. Classes are assigned based on phylogeny and structural features of the RBS, following^17,18^. Black line denotes the isotonic regression fit (*i*.*e*., piecewise constant, monotonous least square fit) using all points until 2010. Error bars indicate standard deviation. A downward trend is observed for over 40 years (Spearman correlation coefficient: −0.82, p = 10^−18^). **B**. ScanNet-predicted protein binding propensity (higher is better) vs antigenicity (lower is better) of the SARS-CoV-2 RBS for WT, four VOCs, all single-point mutants and 1,000 artificial variants with 15 mutations from WT (same number as Omicron) generated using a sequence generative model (Methods). Crosses indicate 95% confidence interval. Both properties are correlated, illustrating the evolutionary trade-off between ACE2 binding and immune escape. Only 7.4% of the artificial variants are strictly better than Omicron (in the sense of higher binding and lower antigenicity, shaded square), confirming that Omicron was selected for high binding and low antigenicity, while also suggesting ample additional space for further decrease.

SARS-CoV-2 was only recently introduced into human populations with insufficient sequence information for evolutionary analysis. To explore the potential space for additional reduction of antigenicity under ACE2 binding constraints, we generated a repertoire of stability-preserving *de novo* RBD variants (a total of 1,000 variants-each contains 15 point mutations similar to those of Omicron) and analyzed their antigenic profiles (**Methods**). Interestingly, our analysis identified multiple variants with significantly lower antigenicity than that of Omicron (**Figure 3B**), implying that future decrease is possible.

## Discussion

In this study, we leveraged computational prediction facilitated by geometric deep-learning (ScanNet) and experimental approaches to systematically investigate the RBD antigenicity. ScanNet provides a rapid means to quantitatively assess the host antibody response against key structures of emerging viruses and their variants. The use of the mouse model for the investigation of RBD antigenicity enables fair comparison among variants, minimizing the potential bias and background complexity that are often associated with clinical samples. Critically, our results provided explicit evidence that Omicron can drastically attenuate its antigenicity across all epitopes on the RBD, which is the key target by neutralizing antibodies. During our manuscript preparation, a preprint reported that an Omicron-specific mRNA vaccine boost appears to provide inferior protection against Omicron infection in non-human primates compared to boost using the WT mRNA vaccine^20^. Moreover, new clinical data suggested that the antibody titers after Omicron breakthrough cases were lower than those of after Delta infection^2–8^. Finally, Omicron convalescent sera from unvaccinated individuals were found to only weakly neutralize Omicron virus while the serum neutralizing activities against other VOCs were below the detection limits^7,21^. Thus, our study is consistent with both the preclinical vaccine trials and clinical convalescent data and provides critical insights into the underlying mechanism of the attenuated host serologic response against Omicron. Cumulatively, our investigations unravel a potential trajectory of future viral evolution and underlie the challenges to develop effective Omicron-specific vaccines.

**Supplementary Figure S1:**
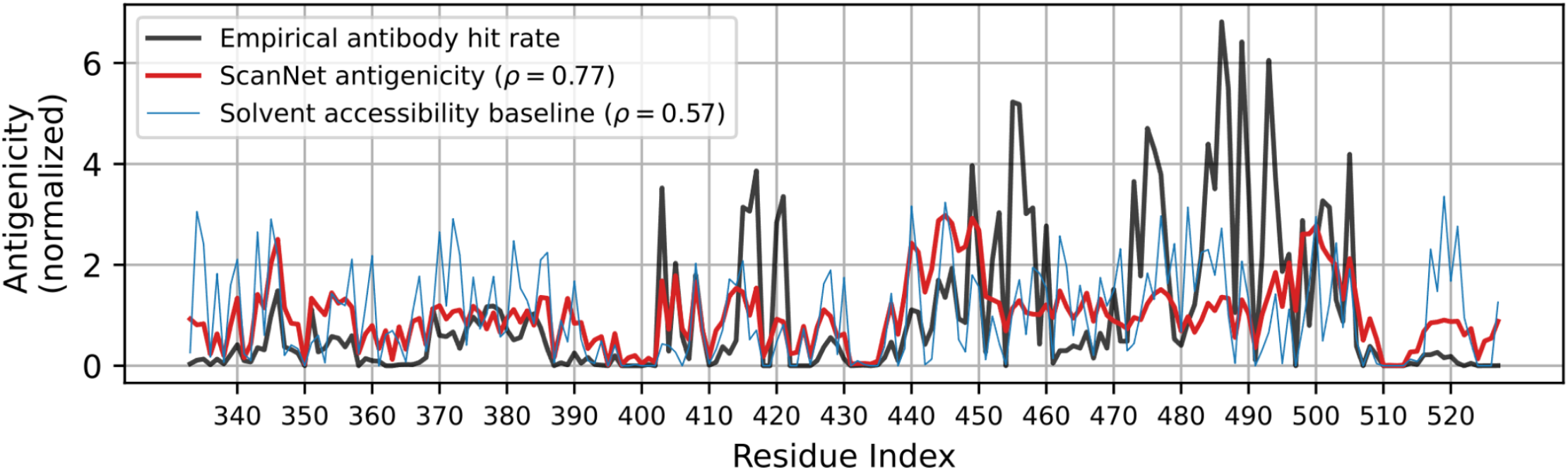
Evaluation of overall accuracy of the ScanNet antigenicity profile on the spike protein RBD. ScanNet antigenicity profile for WT RBD (red) vs. the empirical antibody hit rate calculated based on PDB structures (black). RBD residue solvent accessibility is shown as a baseline (blue). The three curves were normalized to the mean value of 1 to facilitate the comparison. The solvent accessibility was computed within the spike trimer in the open state configuration (PDB identifier: 7e5r:A)

**Supplementary Figure S2.**
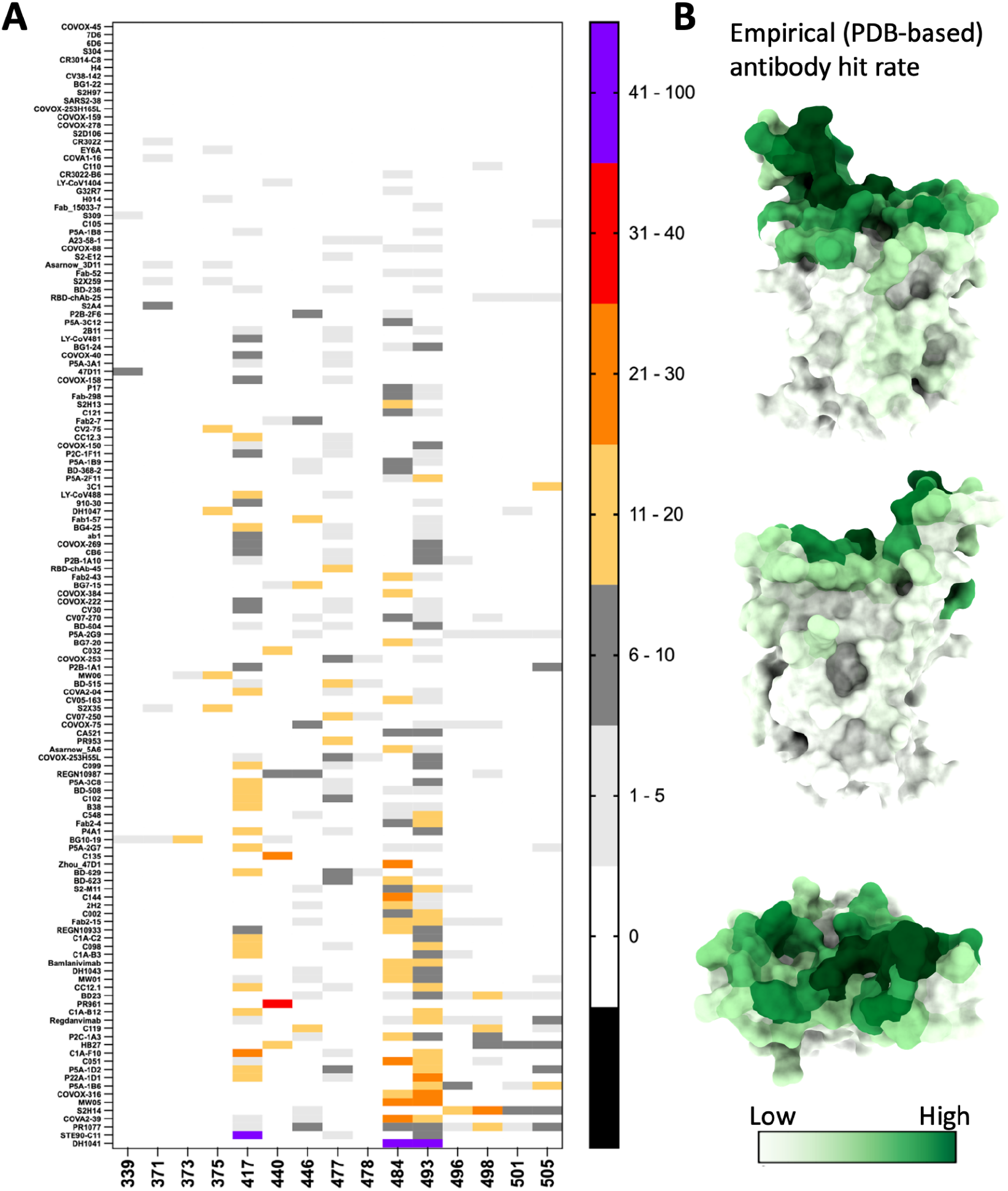
Epitope distribution based on solved antibody-RBD structures. **A**. Omicron mutations (x-axis) vs. antibody binding. The color-coding corresponds to the fraction of the RBD residue atoms interacting with the antibody. **B**. RBD structure (PDB: 7jvb) is colored according to the antibody hit rate (number of antibodies that interact with the residue).

**Supplementary Figure S3:**
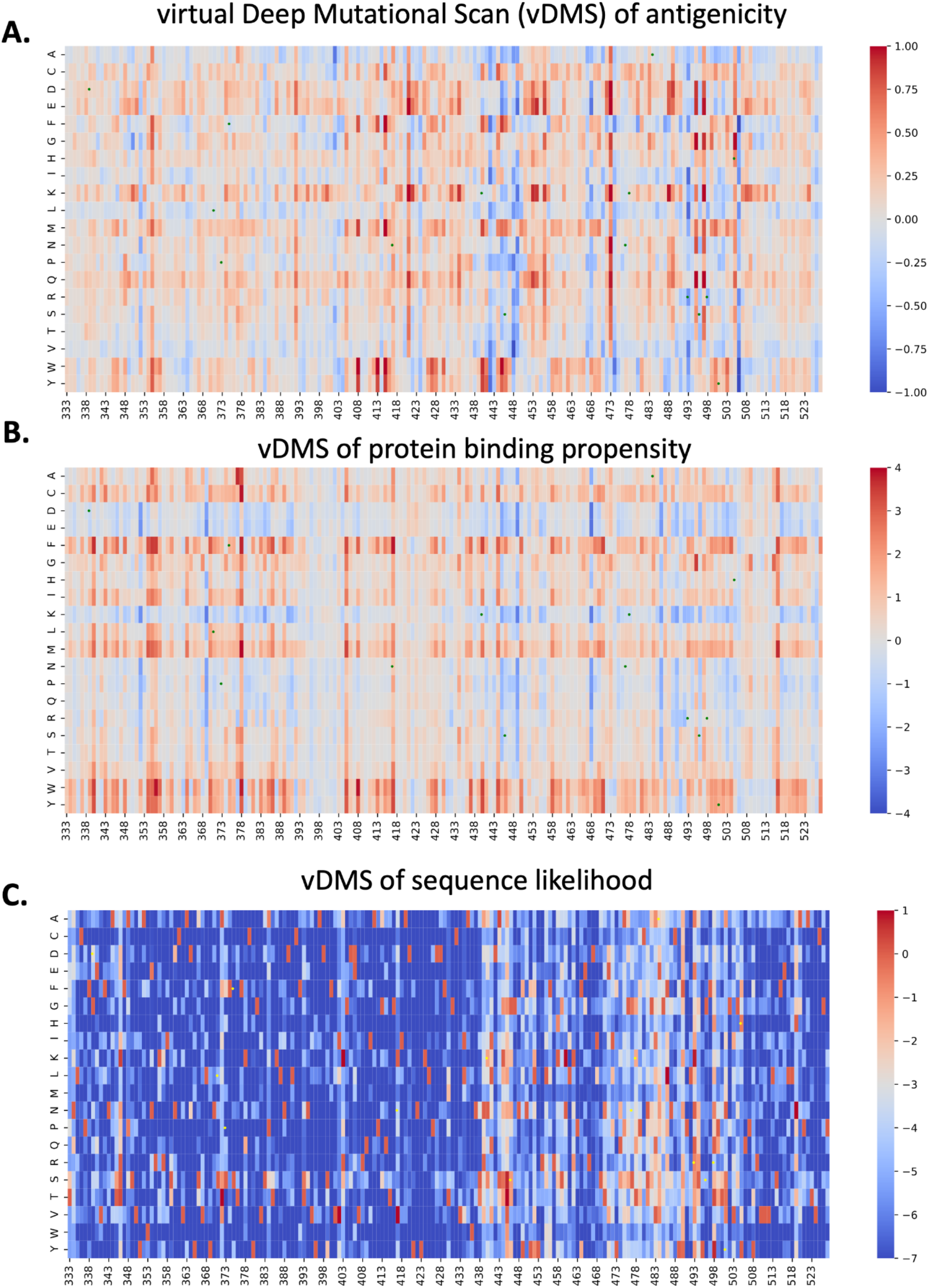
virtual Deep Mutational Scans (vDMS) of WT RBD for antigenicity, protein binding propensity and sequence likelihood. **A**. vDMS of antigenicity score predicted by ScanNet, see Methods. The difference between the *summed* antigenicity profiles (*over the whole domain*) of mutants and WT is shown. The corresponding distribution of entries is shown in **Figure 1F, Supplementary Figure S6B**. Green dots indicate Omicron mutations. **B**. vDMS of protein binding propensity score predicted by ScanNet, see Methods. The difference between the *summed* protein binding propensity profiles *(over the whole domain)* of mutants and WT is shown. The corresponding distribution of entries is shown in **Supplementary Figure S6D**. Green dots indicate Omicron mutations. **C**. vDMS of sequence likelihood based on evolutionary records. The difference between the log-likelihood log P(S) of mutant sequence and wild type sequence is shown. The likelihood function was obtained by training a sequence generative model (Restricted Boltzmann Machine), on a multiple sequence alignment of betacoronaviruses RBDs (**Methods**). The vDMS were averaged over five trainings, each using different random seeds. The correlation to the expression level DMS performed in^22^ is shown in **Supplementary Figure S7D**. Yellow dots indicate Omicron mutations.

**Supplemental Figure S4:**
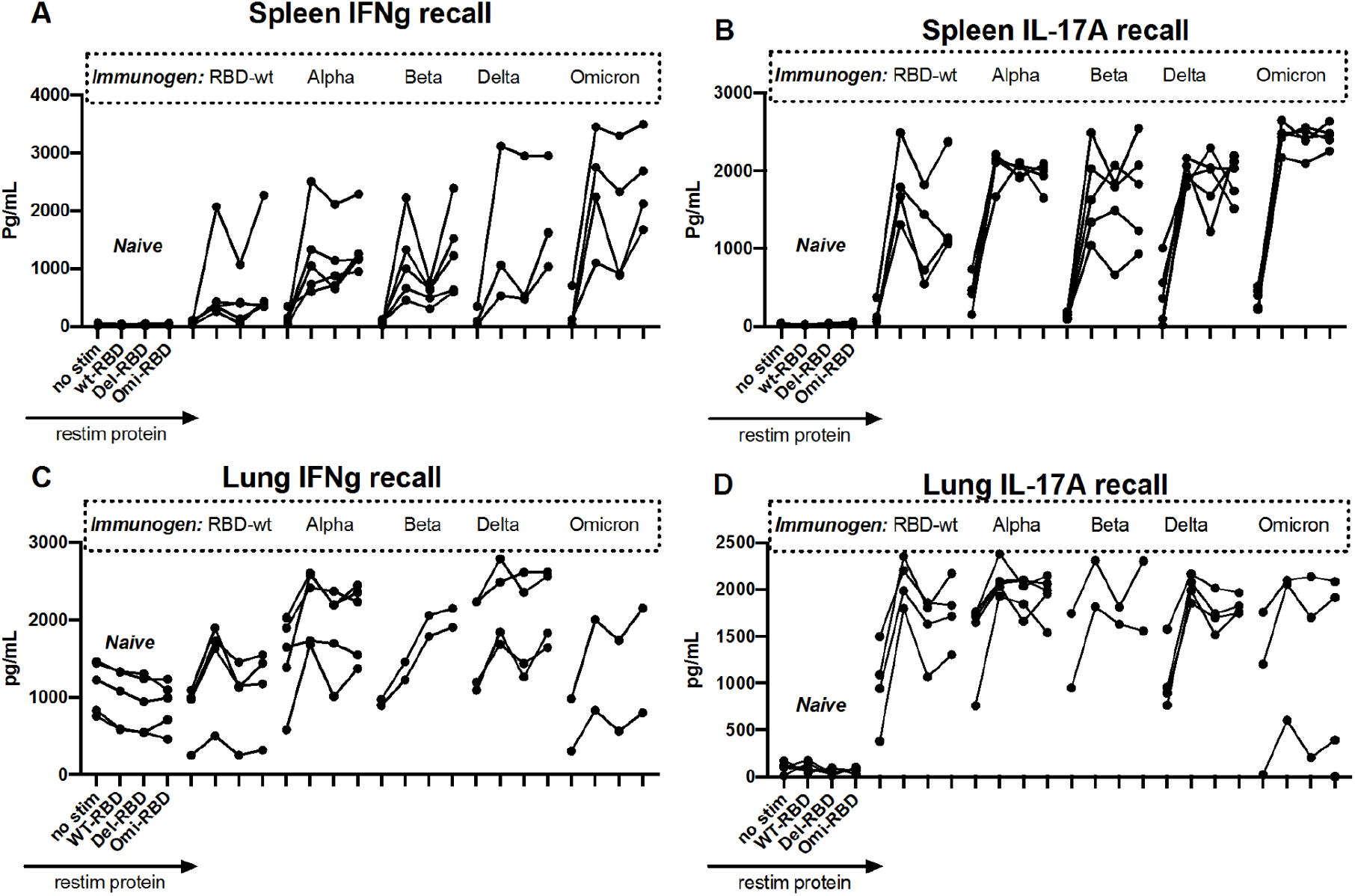
The T cell recall responses in the RBD immunized mice. Splenocytes (**A, B**) or lung mononuclear cells (**C, D**) from naïve or RBD immunized mice were left unstimulated or stimulated with WT, Delta, or Omicron RBD for 72h. Culture supernatants were harvested for IFNg and IL-17 measurements by ELISA.

**Supplemental Figure S5:**
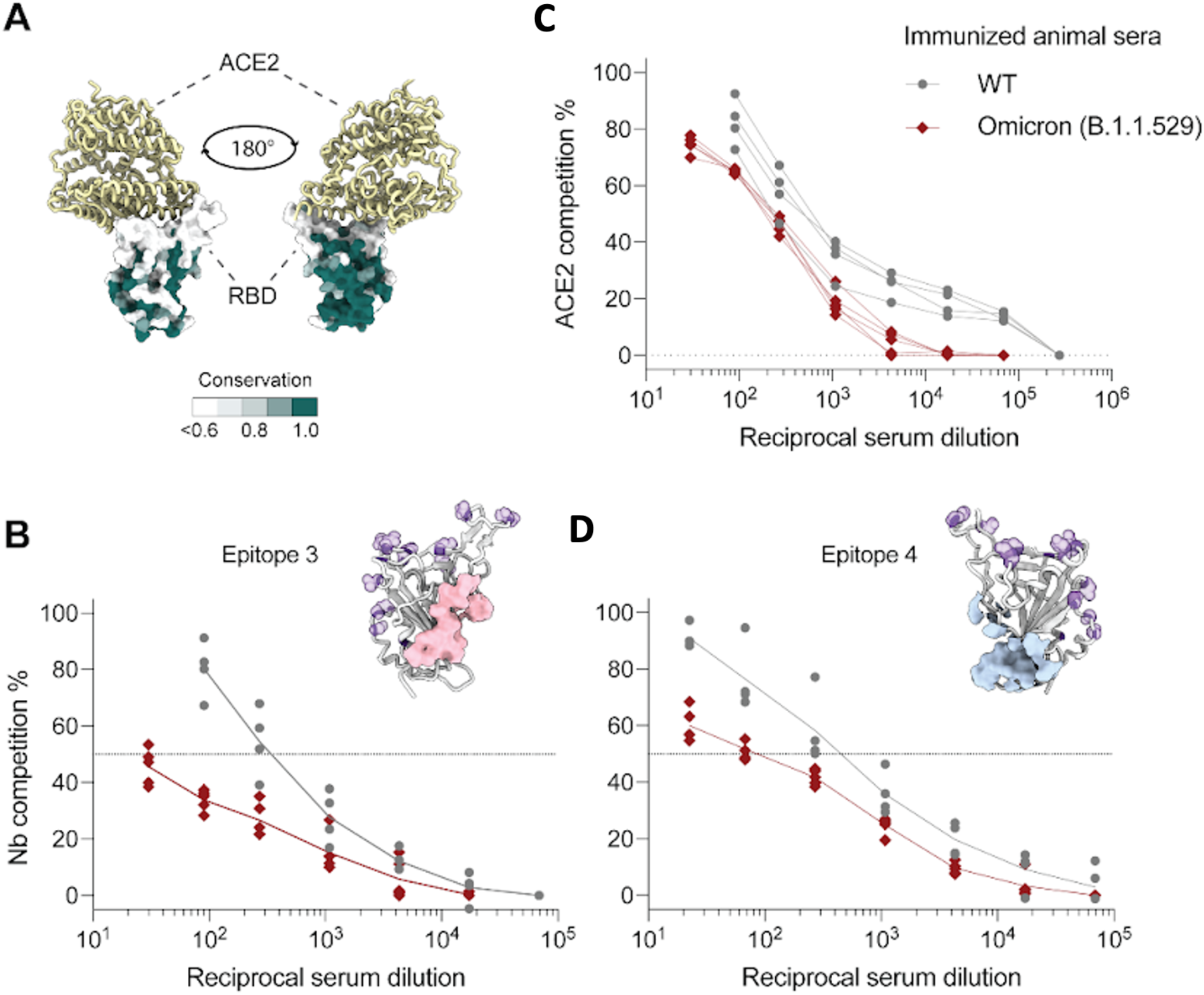
Competition ELISA. **A**. Structural representations of the RBD-ACE2 complex. The sequence conservation of sarbecovirus RBD was presented in a color gradient, where 1.0 (in dark green) indicates that the residue is 100% conserved within all the sarbecovirus clades. **B-D**. Competitive ELISA between mice sera and (**B**) hACE2, (**C**) a high-affinity nanobody that targets a conserved RBD epitope (residues 337, 351-358, 396, 464, 466-468), or (**D**) a high-affinity nanobody that targets another conserved RBD epitope (residues 380, 381, 386, 390, 393, 428-431, 464, 514-522) for RBD binding. Each plot shows the percentage of remaining ACE2 or Nbs on the immobilized RBD in the presence of sera, expressed as reciprocal serum dilution. RBD was shown as gray ribbons. Mutated residues on Omicron were shown in purple. Distinct, conserved nanobody epitopes (3 and 4) were shown in pink and blue, respectively.

**Supplementary Figure S6:**
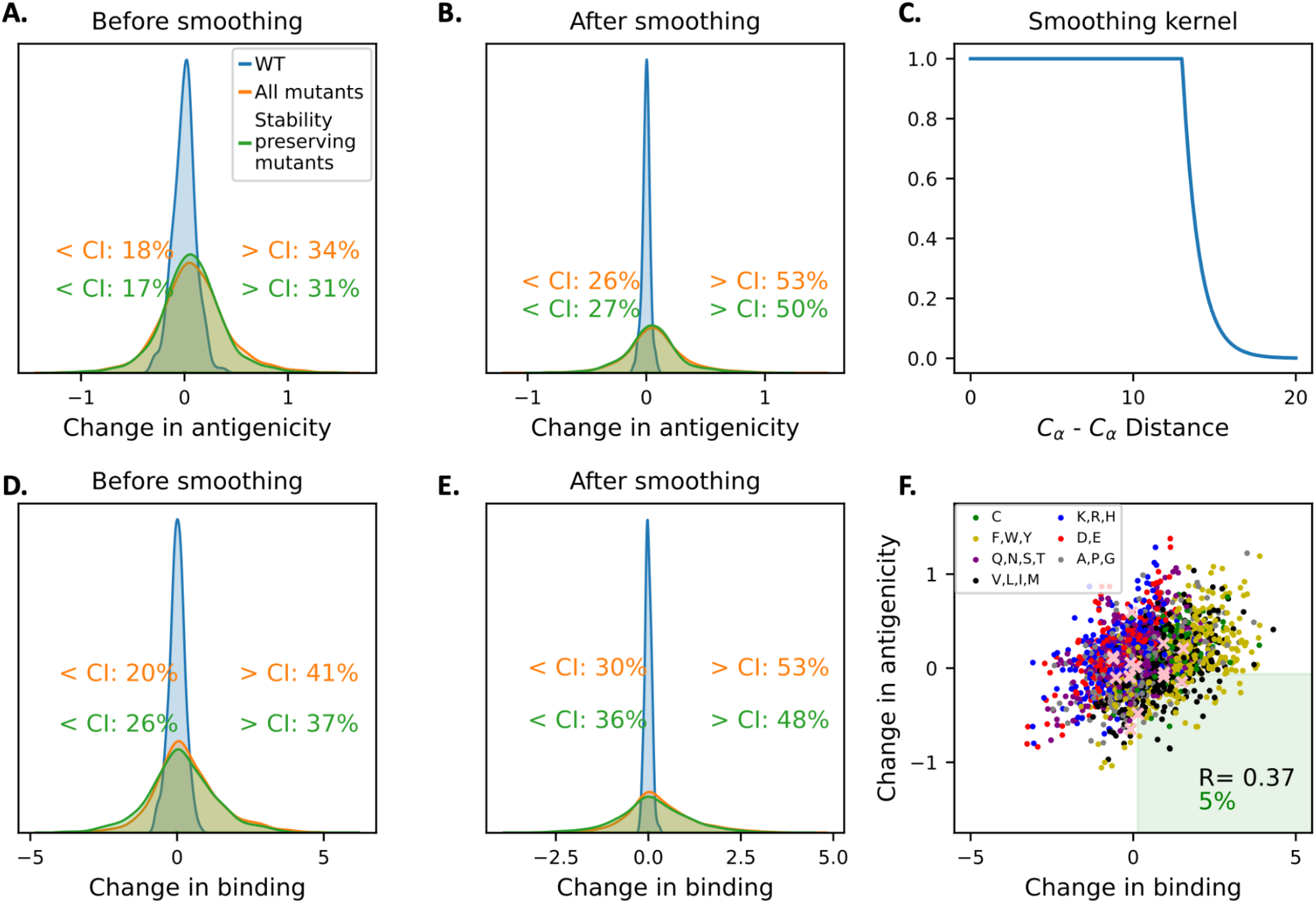
Noise estimation and reduction methodology for assessing the impact of all single point mutations on antigenicity and protein binding propensity. **A-B**. Histogram of the total change in antigenicity with respect to WT across all single point mutations before (**A**) and after (**B**) smoothing. Blue histogram represents 195 repeated runs of the WT sequence through the comparative modeling pipeline; it corresponds to the noise level induced by homology modeling. Text indicates the fraction of mutations outside of the [5%,95%] confidence interval. **C**. The smoothing kernel used for weighting residues in the neighborhood of the mutation. **D-E**. Histogram of the total change in protein binding propensity with respect to WT across all single point mutations before (D) and after (E) smoothing. **F**. Change in protein binding propensity vs. change in antigenicity for all single-point mutants. Each point corresponds to a single mutation, colored by the type of amino acid (green: cysteine, gold: aromatic, purple: polar, black: hydrophobic, blue: positively charged, red: negatively charged, gray: tiny/proline). The green shaded region denotes mutations that simultaneously increase protein binding propensity and decrease antigenicity; they form a small since both properties are correlated. Pink crosses indicate Omicron mutations.

**Supplementary Figure S7:**
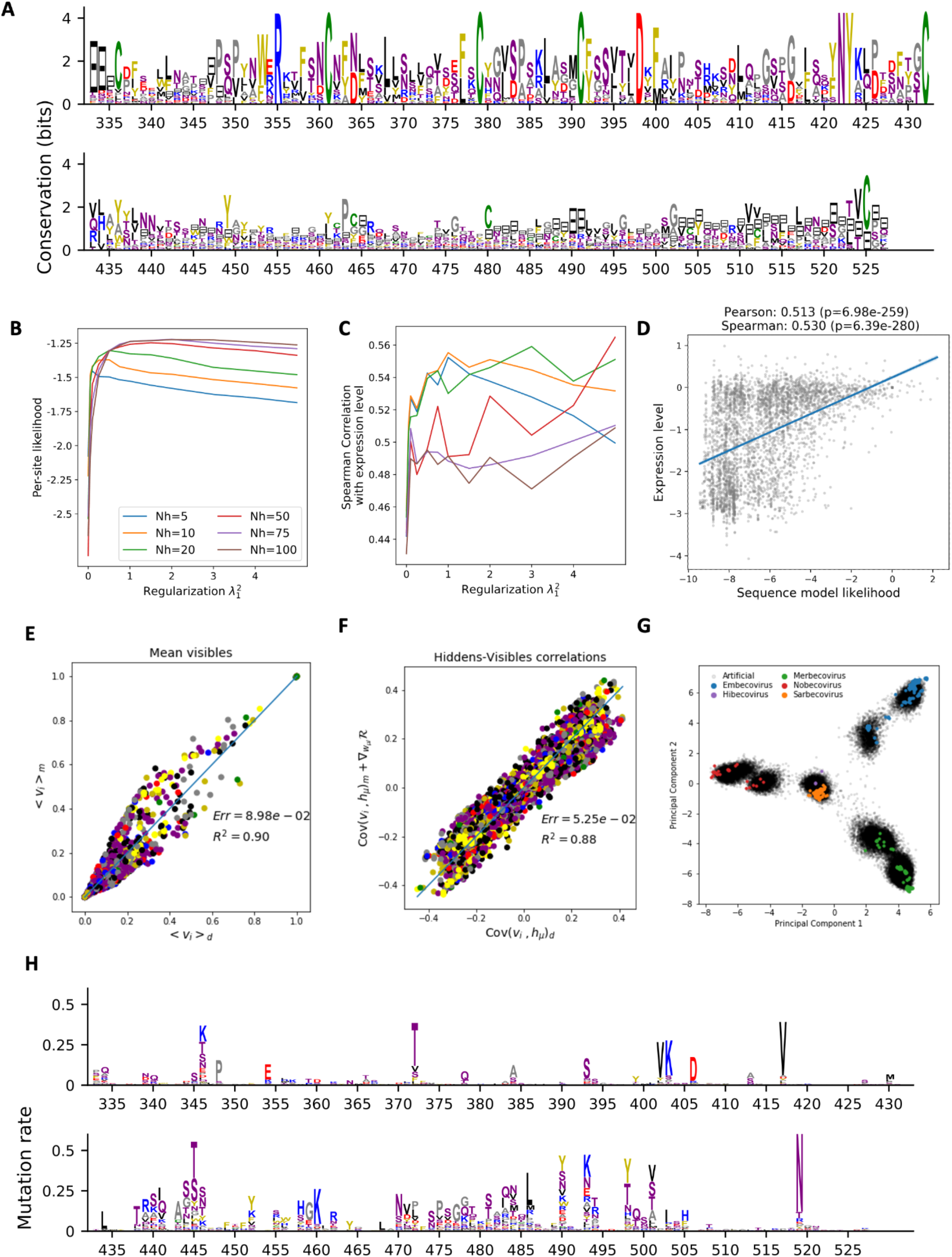
Training, validation and sampling of a sequence generative model for the RBD. **A**. Sequence profile of the MSA of betacoronavirus RBDs identified in UniRef30. **B**,**C**. Hyperparameter search by cross-validation. (**B**) Cross-validation likelihood (divided by the number of sites) and (**C**) Spearman correlation between the change in sequence log-likelihood and change in expression around WT, as function of the number of hidden units and of the regularization strength 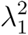 of the RBM. The pseudo-likelihood (not shown) was also monitored and was highly correlated to the likelihood. Since the three metrics did not peak at the same position, we manually selected *M* = 20, 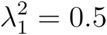 as a compromise. **D**. Scatter plot of the change in expression level (experimentally determined in^22^) and the change in log-likelihood for all single point mutants (corresponding matrix shown in **Figure S3C**). **E-G**. Evaluation of the generative properties of the selected model. The distribution of samples generated by the model matched reasonably well the (**E**) single site frequency, (**F**) covariance between visible and hidden units and (**G**) overall topology of the distribution of natural sequences. **H**. Distribution of mutations to WT found in the set of 1000 artificial variants obtained by sampling from *P*(*S*|*D*_Hamming_ (*S, WT*) = 15, *N*_*gaps*_ = 0. For each site, the height of each letter is proportional to the frequency of the corresponding amino acid in the generated set. The total height is proportional to the mutation rate of the site. As expected, RBS is the most variable region.

## Methods

### ScanNet

Deep learning has been highly successful in protein structure prediction^23–28^. However, leveraging the structures for function prediction has remained a major challenge^29^. Recently, we have developed ScanNet, a geometric deep learning model for structure-based prediction of binding sites including protein-protein binding sites and B-cell epitopes^10^. ScanNet is an end-to-end architecture learning representations of atoms and amino acids based on the spatio-chemical arrangements of their neighbors. Briefly, ScanNet first extracts an atomic neighborhood around each heavy atom (K=16 neighbors, corresponding to about 4Å), and calculates their local coordinates in a frame centered around the atom and oriented using the covalent bonds. The neighborhood, formally a point cloud with attributes (atom group type) is then passed through a set of trainable spatio-chemical filters. Each filter detects a specific spatio-chemical pattern within the neighborhood, such as hydrogen bonds. Conversely, some filters also detect prescribed absences of atoms, e.g. exposed side chain atoms, or backbones nitrogens/oxygens available for hydrogen bond formation. The later filters are critical for epitope prediction, as reactive atom groups that are not engaged in intra-chain interactions are more prone to be targeted by antibodies. The resulting atom-wise embeddings are next pooled at the amino acid level, and the process is reiterated around each amino acid. Finally, the resulting amino acid-wise embeddings are converted to propensity scores via a neighborhood attention module, which projects the embeddings to scalar values and smoothes them (in a learnt fashion) across a neighborhood.

We previously trained ScanNet for detecting B-cell epitopes based on 3756 antibody-antigen complexes available from the PDB. ScanNet predicted known epitopes substantially more accurately than previous works that relied on amino acid propensity scores and geometric features such as solvent accessibility. We previously found that for the Spike protein RBD of WT, the predicted antigenicity profile correlated well with the residue-wise antibody hit rate computed from 246 PDB structures of spike protein - antibody complexes, defined as the fraction of antibodies that bind to the residue^10^. We successfully reproduced the analysis with the prediction pipeline described below (**Figure S1**).

### Antigenicity and protein binding profiles of VOCs

Since the sequences considered are highly similar (92-99% sequence identity to WT) and the structures are virtually indistinguishable by human eye, the predicted epitope propensity profiles are overall similar. Additionally, ScanNet is sensitive to subtle structural features such as sidechain-backbone hydrogen bonds (especially for asparagines) that are not always consistent from one crystal structure to the other for a given variant. To maximize the signal-to-noise ratio, we proceeded as follows:

1. We used the model version that only takes the sequence and structure as input and discards the position-weight matrix. For the antigenicity profile, this version achieves the same performance as the one using evolutionary information (Table S4^10^). For the protein binding profile, the performance is overall lower than the version using evolutionary information (Table 1^10^), but is nonetheless satisfactory for the Spike RBD.
2. All predictions were averaged over 11 networks, each trained using a different random seed. All SARS-CoV-1/2 antibody-antigen complexes were excluded from the training set.
3. We used multiple RBD structures per variant. For the WT, we selected 29 RBD structures. For the other VOCs, all the available RBD structures were taken (WT PDBs: 7eam:A, 7mzj:B, 7dhx:B, 7mfu:A, 7efr:B, 7kn3:A, 7mmo:C, 7kgj:A, 7n4j:A, 7mf1:A, 7mzm:A, 7jmo:A, 7vnb:B, 7s4s:A, 7lop:Z, 7r6w:R, 7kmg:C, 7deu:A, 7det:A, 7c8v:B, 7cjf:C, 7d2z:B, 7bnv:A, 7nx6:E, 6m0j:E, 7mzh:E, 7ean:A, 7n3i:C, 6yla:E. Alpha: 7fdg:E, 7neg:E, 7nx9:E, 7mji:B, 7mjl:A, 7mjn:B, 7ekf:B. Beta: 7ps4:E, 7ps6:E, 7ps0:E, 7ps7:E, 7ps2:G, 7ps0:A, 7ps5:E, 7q0h:E, 7prz:E, 7pry:E, 7ps1:E, 7q0g:E, 7nxa:E, 7e8m:E. Delta: 7w9f:E, 7w9i:E, 7wbq:B, 7wbq:D, 7v8b:A. Omicron: 7qnw:E, 7wbp:B, 7wbl:B, 7t9l:A. SARS-CoV-1: 3bgf:S, 7rks:R, 6waq:D, 2ajf:E, 3d0g:E, 3scl:E, 2ghv:E, 2ghw:A, 2dd8:S).
4. Since some structures consistently missed many sidechain-backbone hydrogen bonds, we standardized them by applying to each structure the FastRelax protocol of PyRosetta (5 cycles)^30,31^. To reduce the noise induced by Rosetta, we generated 20 relaxation runs per structure and averaged epitope profiles over them. This protocol reduced the intra-variant, inter-structure standard deviation by 10-25%.

Altogether, the antigenicity and protein binding propensity profiles of each single point mutant were averaged over 11 × 20 × N profiles where N was the number of available RBD structures. Based on the intra-variant, inter-structure variance, we estimated the average resolution of our differential antigenicity profiles as 0.008 (in probability units).

### Antigenicity and protein binding profiles of all single point mutants

Mutant structures were generated using comparative modeling. We first selected six representative templates for the WT RBD by clustering the aforementioned RBD structures (7jvb:A, 7eam:A, 7d2z:B, 7kgj:A, 7vnb:B, 7det:A). Next, we generated for each single point mutant and each template 20 structural models using MODELLER^32^. As neutral controls, we also generated structural models for the original amino acid at each position (i.e. the WT sequence). Each model was scored using the 11 networks, obtaining 6×20×11=1320 profiles per mutant which were then averaged to yield a single antigenicity profile and a single binding propensity profile. The overall impact of a mutation to antigenicity was defined as the difference between the summed profiles across the entire protein. Despite the averaging, we found that conformational variability yielded changes in total propensity of the same order of magnitude as the one of changes upon single point mutants: 48% of the mutations had an insignificant impact on total antigenicity, *i*.*e*. within the [5%,95%] percentiles of the WT antigenicity distribution (**Figure S6A**). The predicted profiles notably featured small variations in regions far away from the mutation, arising solely because of modeling noise. To improve the signal to noise ratio, we instead computed a weighted sum of the difference of profiles, where the weight is a smoothing function of the distance to the mutated residue (**Figure S6C**). Since ScanNet predictions are based on local neighborhoods and the conformational noise away from the mutation is expected to average out anyway, the local estimator is unbiased and has lower variance. After smoothing, only 21% of mutations were insignificant (**Figure S6B**). Protein binding propensity profiles were calculated in the same manner using comparative models and ScanNet models trained for protein binding site prediction (**Figure S6D-E)**. The ~470k structural models and ~10 million profiles were generated in about ten days using a single computer with 64Gb RAM and an Intel Xeon Phi processor with 56 cores (52ms per profile).

### Generation and screening of stability-preserving artificial RBD variants

Here, we investigate whether additional mutations of the SARS-CoV-2 RBD could further reduce its RBS antigenicity. Although the virtual deep mutational scan readily identifies multiple mutations that could lead to antigenicity reduction (particularly on sites 448,449, 506), it is unclear whether or not they are beneficial for the overall viral fitness. Viral fitness indeed includes multiple factors, such as affinity and specificity of ACE2 binding, structural stability, equilibrium distribution of up and down conformations and corresponding transition times. To this end, we restricted the search space to variants that are likely to arise based on past evolutionary records. This was done in four steps: (1) Construction of a multiple sequence alignment (MSA) of beta coronaviruses RBDs. (2) Selection, training and validation of a sequence generative model, i.e. a probability distribution over RBD sequences P(S). (3) Generation of artificial variants by sampling from the sequence generative model in the vicinity of the original WT. (4) Screening for antigenicity and binding using ScanNet.

#### 1. Construction of the MSA

Homologs of the WT RBD were first searched in the UniprotKB using BLAST. Top hits were manually aligned with MAFFT^33^ (command: mafft —amino —localpair —maxiterate 1000 —op 5 —ep 0), and only the columns not gapped for the WT were kept. Next, additional homologs were searched in the UniRef30 (release 2020/06) using HH-blits^34^. After filtering out hits with unknown residues and/or >25% of gaps, we obtained a (redundant) alignment of *B* = 521 sequences. The MSA covered all the five betacoronaviruses subgenii (sarbecovirus, embecovirus, merbecovirus, nobecovirus, hibecovirus). The effective number of sequences (defined as in^35^, approximately corresponding to the number of 90% sequence identity clusters) was *B*_*eff*_ = 72.8, a relatively low value. The sequence profile of the MSA (**Figure S7A**) features conserved sites (most of which are buried), whereas the RBS region is highly variable.

#### 2. Sequence generative model

Herein, the objective is to learn a probability distribution over the sequence *P*(*S*) space by maximizing the average likelihood ⟨log *P*(*S*) ⟩ of the previously observed viral sequences in the MSA. Intuitively, maximizing the likelihood amounts to assigning high probability values to seen (i.e. evolutionary selected) sequences and low elsewhere (i.e. sequences unexplored or washed away by selection), such that is normalized to 1. The likelihood can therefore be interpreted as a proxy for viral fitness^36^. Importantly, a “smooth” parametric form *P*_0_(*S*) must be chosen to ensure that the model also assigns high probability values to sequences that are close (and presumably evolutionary fit), but unobserved either due to limited sequencing or exploration of the sequence space throughout evolution. Possible choices for the parametric forms include the independent model (*i*.*e*. the position specific sequence model or equivalently, insertion-free HMM profiles), Potts model (i.e. the Boltzmann Machine, BM)^35^ or Restricted Boltzmann Machine (RBM) as well as various deep learning-based models^37,38^. We used RBM here, which is an undirected graphical model that learns the conservation and coevolution patterns of the sequence distribution^39^. In the context of RBD modeling, RBM enjoys two desirable properties over the more thoroughly validated BM model. First, its flexible number of parameters allows better optimization of the bias-variance trade-off. RBM has *NX* (*M* + 1) *X*_*q*_ parameters, where N is the number of columns, M is the (tunable) number of hidden units and *q* = 21 is the number of amino acids (+gap) compared to *N* (*N* − 1) /2_*q*_^2^ + *N*_*q*_ for the Potts model. Our selected model has 100X fewer parameters than a regular Potts model. Second, it is able to model high-order epistasis arising from heterogeneous viral fitness landscapes. Indeed, since different subgenii target different receptors, they are expected to have related but distinct fitness landscapes.

RBM were trained using the PGM package (https://github.com/jertubiana/PGM)^39^ using the Persistent Contrastive Divergence algorithm with the following parameters: number of hidden units: from 5 to 100; hidden unit potential: dReLU; batch size: 100; number of Markov chains: 100; number of Monte Carlo steps between each gradient evaluation: 100; number of gradient updates: 40000; optimizer: ADAM with initial learning rate:, exponentially decaying after 50% of the training to 5. 10^−6^, *β*_1_ =0, *β*_2_ = 0.99, *ε* = 10^−3^. For the regularization, we used a 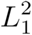 penalty on the weights (of strength 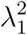 ranging from 0.0 to 5.0) and *L*_2_ penalty on the fields (of strength 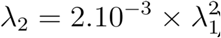). Samples were assigned a weight inversely proportional to their number of 90% sequence identity homologs in the MSA. Annealed importance sampling was used to evaluate the partition functions, using intermediate temperatures and 10 repeats. The low depth of the alignment prompted us to thoroughly explore the hyperparameter space to best calibrate the model complexity (**Figure S7B**,**C**). We divided the MSA into five folds such that any pair of sequences belonging to different folds have at most 80% sequence identity, and performed a grid search over the regularization strength and number of hidden units. We monitored the quality of convergence, the cross-validation likelihood, cross-validation pseudo-likelihood (not shown, correlated to the likelihood), and the spearman correlation between the experimentally determined change of expression level upon mutation (which correlates with structural stability) and the corresponding likelihood difference. We selected the model with *M* = 20 hidden units and regularization strength 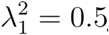. It featured a per-site likelihood value of −1.31 (compared to −1.98 for the best independent model after grid search on pseudo-count values), Spearman correlation *ρ* = 0.53 (**Figure S7D**, compared to 0.42 for the best independent model and 0.54 for the Potts model as recently reported in^40^) and per-site entropy of 0.99 (corresponding to 2.7 amino acid choices per site). Finally, the generative properties of the model were deemed satisfactory: Monte Carlo samples obtained from *P*(*S*) reproduced the moments (**Figure S7E**,**F)** and the clustered topology of the distribution of natural sequences (**Figure S7G)**.

#### 3. Artificial mutant generation

After model selection 1,000 artificial mutants were generated as follows. We sampled from the gap-less, focused distribution *P*(*S*|*D*_*Hamming*_(*WT, S*) = 15, *N*_*gaps*_= 0), where 15 is the same number of mutations from WT as Omicron. Sampling from the conditional distribution was done by importance sampling Markov Chain Monte Carlo, i.e. by sampling from the modified distribution *P*_*λ*_*(S)* ∝ P(S) × exp [*λD*_*Hamming*_*(S,W*) − 10*N*_gaps_(*S*)] where *λ* = 1.9 was chosen such that 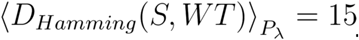. We used the alternate Gibbs sampler, with 5000 burn-in steps, 100 steps between each sample and 100 independent chains. Generated samples with fewer or more mutations were discarded; approximately 5000 samples with exactly 15 mutations were kept. We extracted 1,000 representatives by agglomerative clustering (using as representative the cluster member with highest likelihood). The distribution of the mutations (**Figure S7H**) features high variability on the RBS in general and particularly at positions mutated in VOCs. In total, 1523 of the potential mutations are observed at least once. All VOCs mutations are observed at least once, except for Q498R and S375F, with the mutations Y505H and G446S being the most frequent (in 6.9% and 6.4% of the sequences).

#### 4. Scoring of artificial mutants

We used the same comparative modeling followed by ScanNet antigenicity and binding site prediction pipeline as for the single-point mutants.

We find that 7.4% of the 1000 15-point artificial variants have lower or equal antigenicity and higher or equal binding propensity than Omicron. Such a relatively low percentage is consistent with the hypothesis that Omicron mutations were evolutionary selected for antigenicity reduction under binding constraints. Previous protein-design studies using sequence generative models reported that about 10-80% of artificially generated proteins exhibited native-like functions^38,41–43^. Assuming a conservative estimate of 1% native-like functional sequences, and given that roughly of such variants could be generated, this leaves thousands of potential future variants of concern.

### Longitudinal analysis of the human 229E alphacoronavirus

Previous structural and immunological studies suggested that the hCoV229E alphacoronavirus has been undergoing extensive antibody escape since its entry into human population^17–19^, and that its evolution could reflect the future evolution of SARS-CoV-2^19^. The hCoV229E proteome features a spike protein with a (structurally different) receptor binding domain which targets the human aminopeptidase N protein. Similarly to SARS-CoV-2, the corresponding receptor binding site, which consists of three loops, is also the major immunodominant region. We evaluated the evolution of antigenicity of the hCoV229E RBS using ScanNet as follows. We first collected six template structures for the 229E RBD (PDB 6u7h:A, 6atk:E, 6ixa:A, 6u7e:D, 6u7f:D, 6u7g:D) and constructed a structure-based multiple sequence alignment using ChimeraX^44^, and an HMM profile model (using the hhalign utility^34^). Next, we retrieved all 203 available hCoV229E spike protein sequences from Uniprot, aligned them to the HMM profile (command hhalign -t one_strain_sequence.fasta -i template_sequences.fasta -oa3m output.fasta -all) and discarded sequences that did not cover at least 100 of the 134 columns of the alignment. The corresponding EMBL entry was used to retrieve the corresponding isolate/strain name and its collection date, if available. In total, we obtained 115 (redundant) sequences with known collection dates between 1967 and 2022. For each sequence, we generated 20×6 structural models with MODELLER^32^ (20 per template). Since there was no antibody-hCoV229E spike protein complex in the training set of ScanNet, we used all the 55 networks trained for antibody binding site prediction (including the 11 used elsewhere that were not trained on SARS-CoV-1/2 data). To define the RBS residues, we first extracted all interface residues (6Å distance cut-off) of the template complexes (PDB 6atk, 6u7e, 6u7g, 6u7f) and labeled the corresponding MSA columns as RBS. Then, for a given strain, its RBS residues were identified as the ones mapped onto one of the RBS columns. The RBS antigenicity was defined for a given strain as the average over all networks, all structural models and all non-gapped RBS columns of the antigenicity profile. Note that due to the presence of deletions, the number of residues included in the RBS varied from one variant to the other and therefore summing rather than averaging yielded slightly different results. We tried both options and found a similar decreasing trend in both cases. Error bars (one standard deviation) were estimated based on the structural model variability (**Figure 3A**). The isotonic regression fit was performed using scikit-learn (sklearn.isotonic.IsotonicRegression, default parameters)^32^.

### Mice

8 weeks old female C57BL/6 mice were ordered from The Jackson Laboratory and housed in pathogen-free conditions at the core animal facility at the University of Pittsburgh Medical Center with the approval from the University of Pittsburgh Institutional Animal Care and Use Committee. 40µg recombinant RBD plus 5µg LPS-EB VacciGrade™ (InvivoGen) was given to isoflurane anesthetized mice in sterile PBS (50µL) intranasally on day 0 and day 7 and sacrificed 2 weeks after the last immunization for spleen and lung harvesting.

### *In vitro* Antigen Restimulation Assay

Individual lungs were collected, mechanically digested, and enzymatically digested with collagenase/DNase for 1 hr at 37°C as described previously^45^. Single cell suspensions were then passed through a 70-μm sterile filter. Red blood cells were lysed using a NH4Cl solution and the cells were enumerated then plated at 5 × 10^5^ cells per well in 96-well, stimulated with 10µg/mL recombinant RBD proteins for 72 h. The supernatants were collected and analyzed by murine IFNg and IL-17A ELISA (BioLegend). Spleens were processed similar to the lungs without the need of enzymatic digestion.

### ELISA (enzyme-linked immunosorbent assay)

Indirect ELISA was carried out to evaluate the serological responses of the total antibody in mice sera to an RBD. A 96-well ELISA plate (R&D system) was coated with the recombinant RBD protein (Acro Biosystems) at an amount of approximately 2-3 ng per well in a coating buffer (15 mM sodium carbonate, 35 mM sodium bicarbonate, pH 9.6) overnight at 4°C, with subsequent blockage with a blocking buffer (DPBS, v/v 0.05% Tween 20, 5% milk) at room temperature for 2 hours. To test the immune response, the mice serum was serially 4 or 5-fold diluted starting from 1:27 (Omicron-immunized sera), 1:72 (WT) or 1:100 (other VOCs) in the blocking buffer and then incubated with the RBD-coated wells at room temperature for 2 hours. HRP-conjugated secondary goat anti-mouse IgG (H+L) (Thermo Fisher, cat# G-21040) were diluted 1:1,500 in the blocking buffer and incubated with each well for an additional 1 hour at room temperature. Three washes with 1x PBST (DPBS, v/v 0.05% Tween 20) were carried out to remove nonspecific absorbances between each incubation. After the final wash, the samples were further incubated in the dark with freshly prepared w3,3′,5,5′-Tetramethylbenzidine (TMB) substrate for 10 mins at room temperature to develop the signals. After the STOP solution (R&D system), the plates were read at multiple wavelengths (450 nm and 550 nm) on a plate reader (Multiskan GO, Thermo Fisher). The raw data were processed by Prism 9 (GraphPad) to fit into a 4PL curve and to calculate IC_50_/logIC_50_.

### Competitive ELISA with recombinant hACE2

A 96-well plate was pre-coated with either WT or Omicron recombinant RBD at 2-3 µg/ml at 4°C overnight. Mice serum was 3-fold diluted starting from 1:15 (Omicron) or 1:45 (WT) in the blocking buffer with a final amount of 50 ng biotinylated hACE2 (Sino Biological, cat# 10108-H08H-B) / 8 ng epitope 3 nanobody / 8 ng epitope 4 nanobody at each concentration and then incubated with the plate at room temperature for 2 hrs. The plate was washed by the washing buffer to remove the unbound hACE2. 1:5,000 diluted Pierce™ High Sensitivity NeutrAvidin™-HRP (Thermo Fisher cat# 31030) or 1:7,500 diluted T7-tag polyclonal antibody-HRP (Thermo Fisher, cat# PA1-31449) were incubated with the plate for 1 hr at room temperature. TMB solution was added to react with the HRP conjugates for 10 mins. The reaction was then stopped by the Stop Solution. The signal corresponding to the amount of the bound hACE2 or nanobodies was measured by a plate reader at 450 nm and 550 nm. The wells without sera were used as control to calculate the percentage of hACE2 or nanobody signal. The resulting data were analyzed by Prism 9 (GraphPad) and plotted.

### Pseudotyped SARS-CoV-2 neutralization assay

The 293T-hsACE2 stable cell line (Integral Molecular, cat# C-HA101, Lot# TA060720MC) and pseudotyped SARS-CoV-2 (Wuhan-Hu-1 strain D614G and Omicron) particles with luciferase reporters were purchased from the Integral Molecular. The neutralization assay was carried out according to the manufacturers’ protocols. In brief, 2-fold serially diluted immunized mice serum starting from 1:22 dilution was incubated with the pseudotyped SARS-CoV-2-luciferase. For accurate measurements, seven concentrations were tested for each mice and at least two repeats were done. Pseudovirus in culture media without sera was used as a negative control. 100 µl of the mixtures were then incubated with 100 µl 293T-hsACE2 cells at 2.5×10e^5^ cells/ml in the 96-well plates. The infection took ~72 hrs at 37 °C with 5% CO_2_. The luciferase signal was measured using the *Renilla*-Glo luciferase assay system (Promega, cat# E2720) with the luminometer at 1 ms integration time. The obtained relative luminescence signals (RLU) from the negative control wells were normalized and used to calculate the neutralization percentage at each concentration. Data was processed by Prism 9 (GraphPad). Due to the poor neutralization of the serum at the highest concentration (lowest dilution), the IC_50_ was estimated as the maximal dilution that could inhibit ~50% cell infections by the pseudovirus.

## Acknowledgments

We thank Zhe Sang for the analysis of antibody binding. J.T. acknowledges helpful discussion with Andrea Di Gioacchino and Simona Cocco. **Funding:** This work was supported by NIH grant R35GM137905 (Y.S.), R01HL137709 (K.C.), ISF 1466/18 and Israeli Ministry of Science and Technology (D.S.), the Edmond J. Safra Center for Bioinformatics at Tel Aviv University and from the Human Frontier Science Program (cross-disciplinary postdoctoral fellowship LT001058/2019-C) (JT), Len Blavatnik and the Blavatnik Family Foundation (H.J.W.). **Author contributions:** D.S. and Y.S. conceived the study. J.T. performed all the computational analysis with the help of D.S. and H.J.W. Y.X., L.F., and K.C. performed the experiments. J.T.,Y.S. and D.S drafted the manuscript with substantial input from Y.X. and K.C. All authors reviewed the manuscript. **Competing interests:** no competing interests. **Data availability:** The datasets generated during and/or analysed during the current study are available from the corresponding author on reasonable request. **Code availability:** Source codes for reproducing the computational analysis will be made available before publication.

